# Fendioxypyracil Exhibits Potent Inhibition of PPO-Resistant Mutant Enzymes and Robust Activity Against PPO-Resistant *Amaranthus*

**DOI:** 10.64898/2026.08.23.746358

**Authors:** Aimone Porri, Jens Lerchl, Ingo Meiners, Liliana Parra Rapado, Scott Asher, Sarah Stilgenbauer, Jason K. Norsworthy, Susee Sudhakar

## Abstract

**Background:** Resistance to protoporphyrinogen oxidase (PPO)-inhibiting herbicides is mainly driven by diverse target-site mutations, reducing the effectiveness of this site of action in row-crop systems. Fendioxypyracil is a newly developed PPO inhibitor with high intrinsic grass and broadleaf activity, but its performance against resistant populations and target-site enzyme variants remains insufficiently characterized.

**Results:** Enzyme assays using PPO2 from *Amaranthus palmeri* and *Setaria viridis* demonstrated that fendioxypyracil maintained low IC_50_ values across a broad range of resistance-associated mutations, including ΔG210 deletion and G210, R128, and G399 substitutions, whereas oxadiazon, tiafenacil, and saflufenacil showed substantial loss of potency. Greenhouse dose– response experiments confirmed strong fendioxypyracil efficacy, with susceptible and G399A populations controlled at <3 g ai ha⁻¹, while ΔG210 and R128G populations showed only moderate shifts in sensitivity but remained effectively controlled at the recommended rate. Transgenic *Arabidopsis thaliana* expressing resistant *PPX2* alleles exhibited faster and more severe injury with fendioxypyracil compared to saflufenacil. Field trials conducted in a PPO-resistant *Amaranthus palmeri* population demonstrated that fendioxypyracil provided consistent weed control and density reduction, matching the performance of trifludimoxazin and saflufenacil while exceeding that of fomesafen.

**Conclusion:** Fendioxypyracil provides robust and broad-spectrum activity against PPO-resistant *Amaranthus* populations and target mutant enzymes, maintaining efficacy across diverse mutation backgrounds. These results demonstrate its potential as an effective tool for managing PPO inhibitor resistance and sustaining weed control in row-crop production systems.

## 1. Introduction

*Amaranthus palmeri (Amaranthus palmeri* S. Watson) is one of the most troublesome weeds in United States row crops because of its rapid growth, extended emergence period, high seed production, and strong ability to evolve herbicide resistance.^1^ It is a problematic weed in soybean and cotton production systems, where repeated use of the same herbicide sites of action has selected for resistance to several herbicide groups.^2^ The widespread occurrence of resistance to acetolactate synthase (ALS) inhibitors and glyphosate in *Amaranthus palmeri* in row-crop systems led to the use of protoporphyrinogen oxidase (PPO)-inhibiting herbicides in postemergence programs and overlapping residual systems.^3,4^ PPO inhibitors prevent the oxidation of protoporphyrinogen IX to protoporphyrin IX, a key step in tetrapyrrole biosynthesis required for chlorophyll and heme formation. In susceptible plants, inhibition of this pathway leads to the accumulation of photodynamic intermediates, membrane disruption, and the rapid leaf necrosis associated with this herbicide group.^5^ Although PPO1 was long considered the primary herbicide target, more recent work has shown that resistance in *Amaranthus* is associated largely with mutations in the *PPX2* gene.^6^

Widespread use of PPO inhibitors in row-crop systems in recent decades has led to the emergence of resistance in several *Amaranthus* species. Resistance to these herbicides was recognized later in *Amaranthus palmeri* than in *Amaranthus tuberculatus*. *Amaranthus palmeri* is naturally more tolerant to postemergence (POST)-applied PPO inhibitors than *Amaranthus tuberculatus*, especially at larger growth stages.^7^ The first confirmed PPO-resistant *Amaranthus palmeri* population in Arkansas was reported in Lawrence County, where fomesafen resistance frequency increased from 5% in the original population to 17% after two cycles of field selection.^3^ This population showed 6-fold to 21-fold greater tolerance in the mean response relative to the sensitive standard, and surviving plants carried the ΔG210 deletion in *PPX2*. In Arkansas, 167 of 227 accessions collected in 2016 from 29 counties showed reduced response to fomesafen treatment, indicating that resistance was already widespread in a major production region.^8^ These populations contained the following mutations: ΔG210 and R128G or R128M. Later studies identified G399A and V361A as additional mutations involved in herbicide resistance.^9,10^ A Mid-South USA survey also showed that some populations carry stacked mutations such as ΔG210 and G399A.^11^ Moreover, double *PPX2* mutations can occur in the same plant, but they are usually found in the heterozygous state because plants with homozygous double mutations are generally not viable.^12^ These studies show that PPO inhibitor resistance in *Amaranthus palmeri* is genetically diverse.

Additionally, populations with no target-site mutations have been found to be resistant to PPO inhibitors. A study in Arkansas showed that pretreatment with malathion or NBD-Cl reduced survival and biomass of a fomesafen-resistant accession, suggesting P450- and GST-based metabolism. ^13^ Another study in Kansas demonstrated rapid metabolism of fomesafen.^14^ Variations in metabolic rate also contribute to differences in cross-resistance among PPO inhibitors.^15^ Resistance level can also vary with genotype; plants harboring the homozygous ΔG210 mutation, particularly those carrying both ΔG210 and G399A, exhibited stronger cross-resistance.^16^

The increasing complexity of resistance has led to interest in newer PPO inhibitors. Among these, trifludimoxazin and epyrifenacil have been extensively researched and generally have shown better activity than older PPO-inhibiting herbicides across *Amaranthus palmeri* populations.^17–19,24^ Trifludimoxazin was developed to maintain activity in plants harboring *PPX2* mutations. Greenhouse evaluations showed that plants carrying either the ΔG210 or V361A mutation alone were sensitive to trifludimoxazin.^17,18^ Epyrifenacil performed well across many accessions, although some ΔG210 populations showed lower sensitivity.^19^ Further, reduced sensitivity to trifludimoxazin has been reported in a resistant population from the state of Georgia.^20^

Fendioxypyracil is a recently developed herbicide intended to expand POST weed-control options. Information on this herbicide is still limited, but available reports describe it as a systemic PPO inhibitor with strong activity against both PPO1 and PPO2 and greater enzyme-level potency than saflufenacil in *Amaranthus* assays. Strong control of *Amaranthus palmeri* at 16 g ai ha⁻¹ has also been reported in greenhouse studies.^21^ However, its evaluation in resistant populations, including enzyme assays, remains limited, and the resistance profile of fendioxypyracil in resistant *Amaranthus palmeri* populations has not yet been fully described. *Amaranthus palmeri* populations now vary widely in PPO response because of ΔG210, R128 substitutions, G399A, V361A, mutation stacking, and non-target-site metabolism. Older PPO inhibitors can be strongly affected by these resistance mechanisms, and even newer chemistries show some limitations depending on the resistance background. Therefore, the objectives of this study were to (i) characterize the inhibitory potency of fendioxypyracil against wild-type and PPO2 target-site mutant enzymes associated with PPO-inhibitor resistance, (ii) evaluate its whole-plant efficacy in susceptible and resistant *Amaranthus* biotypes under greenhouse conditions, (iii) assess herbicide response in transgenic *Arabidopsis* expressing resistant PPO2 alleles, and (iv) determine its field performance relative to commercial PPO herbicides in resistant *Amaranthus palmeri* populations.

## 2. Materials and Methods

### 2.1 Plant material and growth conditions

Seeds were provided by BASF which included three PPO-inhibitor resistant populations. The populations comprised one *Amaranthus tuberculatus* population, ΔG210-biotype, and two *Amaranthus palmeri* populations G399A-biotype substitution and R128G-biotype substitution. A *Amaranthus palmeri* susceptible standard was included for comparison. Seeds were sown in a greenhouse at the Milo J. Shult Agricultural Research and Extension Center. Plants were grown in a commercial potting mix (Pro-Mix® LP15, Premier Horticulture, PA, USA) under a 16-h photoperiod provided by light-emitting diode (LED) lighting, with day/night temperatures of 35/25 °C. Seedlings were transplanted at the 1-leaf stage into 50-cell plastic trays for further growth.

### 2.2 Fendioxypyracil dose–response assay in greenhouse conditions

Plants were treated with fendioxypyracil (BASD) at the 4-to 5-leaf stage. The herbicide was applied at 0.125X, 0.25X, 0.5X, 1X, 2X, and 4X rates, where 1X corresponded to 25 g ai ha⁻¹. All treatments included 1% v/v methylated seed oil (MSO) as an adjuvant. Herbicide applications were performed in a spray chamber equipped with a two-nozzle boom fitted with 1100067 flat-fan nozzles (TeeJet® Technologies, Springfield, IL, USA), calibrated to deliver 187 L ha⁻¹ at 1.6 km h⁻¹. The experiment was arranged in a completely randomized design. In the first two experimental runs, 25 replicates were included per treatment rate, except for the 0.5X and 1X rates, which included 50 replicates. In the third run, replication varied by population: G399A-biotype included 50 replicates per rate; R128G-biotype included 25 replicates per rate; and ΔG210-biotype included 15 replicates at 0.125X, 20 replicates at 0.25X, and 25 replicates for the remaining rates. Each replicate consisted of five plants. The dose–response assay was conducted three times. Following treatment, plants were maintained under greenhouse conditions for further evaluation. At 21 days after treatment (DAT), visible injury was assessed on a scale of 0% to 100% (0% = no injury, 100% = plant death) based on chlorosis and growth inhibition. Plant survival was recorded at the time of evaluation. Aboveground biomass was harvested at 21 DAT, dried at 66 °C for 5 days to constant mass, and weighed. Biomass was expressed relative to the nontreated control for each population.

### 2.3 Fendioxypyracil response in field conditions

A field experiment was conducted in 2025 at Fayetteville, Arkansas, where a previously described and characterized *Amaranthus palmeri* population corresponding to CCR was established and maintained. In a previous study, this population was characterized as less sensitive to fomesafen.^22^ Experiments were arranged in a randomized complete block design with four replications. Plot size was 1.8 × 4.5 m. Herbicide treatments included fendioxypyracil applied at 12.5, 18.75, 25, and 37.5 g ai ha⁻¹, along with commercial standards including fomesafen (Reflex®) at 420 g ai ha⁻¹, saflufenacil (Sharpen®) at 25 g ai ha⁻¹, and trifludimoxazin at 25 g ai ha⁻¹. All postemergence (POST) applications were made with methylated seed oil (MSO) at 1% v/v. Applications were made to two *Amaranthus palmeri* growth stages, targeting plants approximately 5 cm (early POST) and 12.5 cm tall (late POST). Herbicides were applied using a CO₂-pressurized backpack sprayer calibrated to deliver 143 L ha⁻¹ at 4.8 kph. *Amaranthus palmeri* control was visually estimated at 28 days after treatment (DAT) using a scale of 0 to 100%, where 0 indicated no control and 100 indicated complete plant death. Plant density was assessed at 28 DAT for CCR. The sequencing results for the CCR population is represented in table 4.

### 2.4 Statistical analysis

All analyses were conducted in R v4.5.1 (R Core Team 2026). Dose–response data were analyzed separately for each endpoint (growth reduction [GR], visible control [ID], and lethality [LD]) and population. Experimental runs were pooled at the rate × population level, and cell means with associated standard errors were used for modeling. Dose–response curves were fitted using a four-parameter log-logistic model (LL.4) implemented in the *drc* package in R. The model estimates the slope (*b*), lower asymptote (*c*), upper asymptote (*d*), and the effective dose producing a specified level of response (*e*), and is defined as:

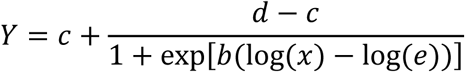

For all endpoints, the lower and upper asymptotes were fixed at 0 and 100, respectively, to constrain responses within biologically meaningful limits. A small positive constant (epsilon) was substituted for the zero rate to enable model fitting on the log scale. The effective dose required to achieve 90% response (ED_90_) was estimated using the (ED) function in *drc*, and standard errors were obtained using the delta method. Dose values were initially modeled on a relative scale (X-rate) and subsequently converted to absolute units (g ai ha⁻¹) using the recommended field rate (1X = 25 g ai ha⁻¹).

Control and density reduction data from the field trials were analyzed using generalized linear models separately for each site. Fixed effects included herbicide treatment, weed size (2-inch and 5-inch *Amaranthus palmeri*), and their interaction. Type III analysis of deviance was performed using Wald χ² tests with the Anova() function in the *car* package to evaluate the significance of main effects and interactions. For density data, counts of *Amaranthus palmeri* plants m⁻² were used for analysis. Significance was determined at α = 0.05.

### 2.5 Evaluation of median inhibitory concentration (IC50)

Complete description of the expression and purification of PPO2 enzyme variants (from *Amaranthus palmeri* and *Setaria viridis*) and the enzymatic assay to determine protein activity and median inhibitory concentration (IC_50_) is provided in Rangani et al.^9^ For the *Setaria* PPO2 enzyme, only catalytically active variants at residues A213, R128, and G409 were included in the analysis. Consequently, the experimental dataset comprised fewer than the 19 possible amino acid substitutions at each residue.

### 2.6 *Arabidopsis* transgenics growth and herbicide treatment

*Arabidopsis thaliana* (accession MC24) seeds were sown in GS90 substrate supplemented with 5% sand, stratified for 5 days at 4 °C, and initially grown for 10 days under short-day conditions (10 h light/14 h dark, 20/18 ± 1 °C, ∼120 μmol m⁻² s⁻¹ PAR). Seedlings were transplanted into pots containing GS90 soil and maintained for a further 14 days before being transferred to long-day conditions (16 h light/8 h dark, ∼200 μmol m⁻² s⁻¹ PAR). Plants were fertilized twice weekly with 0.3% Hakaphos Blau, and relative humidity ranged between approximately 40% and 70%.

Wild-type and mutant *PPX2* constructs from *Amaranthus tuberculatus* were introduced into Agrobacterium tumefaciens and used to generate transgenic plants via the floral dip method. An ALS resistance marker enabled selection of transformants using imazamox (20 ppm). Overexpression lines corresponding to the wildtype *PPX2*, and *PPX2* encoding ΔG210 deletion, G399A and R128G substitutions were selected for herbicide evaluation.

Selected plants were transplanted and grown to the rosette stage (approximately 10 leaves) prior to treatment. Herbicides were applied as foliar sprays using a calibrated spray chamber delivering 375 L ha⁻¹, with DASH HC included as adjuvant. Fendioxypyracil and saflufenacil were applied at 80, 30, 10, and 5 g ai ha⁻¹ in lines expressing mutant *PPX2* allele encoding target site variants, whereas plants expressing wild-type *PPX2* were treated at 30, 10, 5, and 1 g ai ha⁻¹ to capture differential sensitivity. Herbicide response was visually assessed 7 days after application based on the extent of chlorosis/bleaching, inhibition of new leaf development, and tissue necrosis relative to untreated controls.

## 3. Results

### 3.1 Fendioxypyracil resistance profile against Amaranth ΔG210, R128, and G399 target-site mutant enzymes

The inhibitory potency of five PPO-inhibiting herbicides—fendioxypyracil, epyriflenacil, tiafenacil, oxadiazon, and saflufenacil—was evaluated using the *Amaranthus palmeri* wild type and most prevalent target mutant PPO2 enzymes. IC₅₀ values revealed clear differences in PPO2 sensitivity between the wild type and resistance-associated variants (Table 1A). The wild-type enzyme displayed uniformly high sensitivity across all herbicides, with IC₅₀ values in the low nanomolar range and full enzymatic activity.

**Table 1A.** Comparative IC₅₀ values of *Amaranthus palmeri* PPO2 wild type and target site enzyme variants. Half-maximal inhibitory concentrations (IC₅₀, M) of fendiopyracil, epyrifenacil, tiafenacil, oxadiazon, and saflufenacil are shown for wild-type PPO2 and the variants ΔG210, R128G, and G399A. Values are presented in scientific notation and visualized as a heat map, where green indicates low IC₅₀ values (high sensitivity), yellow/orange represents intermediate inhibition, and red indicates high IC₅₀ values (reduced sensitivity). The remaining protein activity for each variant is given as a percentage in the final column.

| Enzyme | Variant (AMAPA) | Fendioxypyracil | Epyrifenacil | Tiafenacil | Oxadiazon | Saflufenacil | Remaining protein activity % |
| --- | --- | --- | --- | --- | --- | --- | --- |
| PPO2 | Wild type | 4.79E-10 | 2.11E-10 | 6.27E-10 | 5.89E-09 | 7.76E-10 | 100 |
| PPO2 | ΔG210 | 1.14E-07 | 2.56E-07 | 1.43E-07 | 1.09E-06 | 2.04E-06 | 10 |
| PPO2 | R128G | 4.20E-09 | 1.59E-09 | 4.56E-09 | 8.57E-09 | 6.61E-07 | 60 |
| PPO2 | G399A | 2.58E-08 | 6.87E-08 | 7.30E-08 | 5.89E-07 | 6.27E-07 | 15 |

The ΔG210 mutation caused the strongest reduction in sensitivity, with IC₅₀ values increasing by several orders of magnitude, particularly for oxadiazon (1.09 × 10⁻⁶ M) and saflufenacil (2.04 × 10⁻⁶ M), and was associated with low enzyme activity (10%). The G399A variant showed an intermediate shift, with consistent increases in IC₅₀ values across all herbicides and reduced activity (15%). In contrast, R128G resulted in only minor changes for most compounds but reduced sensitivity to saflufenacil, while retaining moderate activity (60%).

Across all variants, fendioxypyracil consistently showed the lowest IC₅₀ values indicating superior potency and greater tolerance to target-site mutations. Epyrifenacil and tiafenacil similarly retained high activity across most variants, although an increases in IC₅₀ were observed for ΔG210 and less pronounced for G399A. In contrast, oxadiazon and saflufenacil were strongly affected by the mutations. Together, these results highlight the robustness of fendioxypyracil, with epyrifenacil and tiafenacil showing a similar but slightly less pronounced tolerance to PPO2 target site mutations, especially to the ΔG210.

### 3.2 Fendioxypyracil dose-response under greenhouse conditions

Dose–response analysis indicated that fendioxypyracil maintained strong overall efficacy across the tested populations, with most populations exhibiting high sensitivity at the recommended field rate. The *Amaranthus palmeri* population G399A -biotype and the susceptible standard S2 showed GR90, ID90, and LD90 values below the lowest tested rate (<3 g ai ha⁻¹), indicating excellent control. Visual assessments supported these findings; with rapid injury development and complete mortality observed at the recommended field rate (Table 2; Fig. 2).

**Fig 2.**
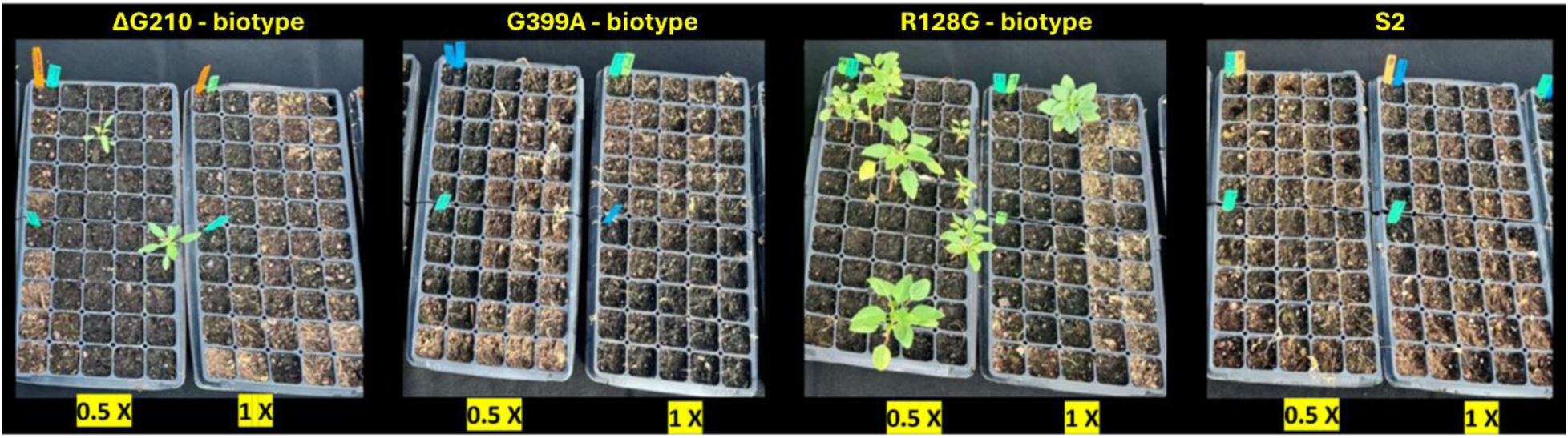
Visual response of *Amaranthus palmeri and tuberculatus* populations carrying different *PPX2* target-site mutations (ΔG210-biotype, G399A-biotype, and R128G-biotype) and the sensitive population (S2) following application of fendioxypyracil at 0.5× and 1× of the recommended rate in greenhouse condition. Photographs were taken 21 days after treatment.

The *Amaranthus tuberculatus* population ΔG210-biotype exhibited a moderate shift in response relative to the susceptible population, with GR_9_₀, ID_90_, and LD_90_ values of 3.74, 5.97, and 6.95 g ai ha⁻¹, respectively (Table 2; Fig 2). However, visual observations demonstrated a full control achieved at 1X. Similarly, the *Amaranthus palmeri* population R128G-biotype showed a comparatively reduced response compared to the susceptible population with GR_90_, ID_90_, and LD_90_ values of 5.0, 12.10, and 13.23 g ai ha⁻¹, respectively, but with substantial control observed at the recommended field rate.

Thus, across all populations harboring distinct PPO2 amino acid substitutions, fendioxypyracil provided effective control at the recommended field rate. Although differences in response were observed among mutation backgrounds, these variations did not translate into a meaningful reduction in herbicide performance under the conditions evaluated.

### 3.3 Fendioxypyracil efficacy against transgenic Arabidopsis carrying PPO2 target-site mutations

Across all PPO2 variants, fendioxypyracil induces stronger, faster developing, and more uniform injury symptoms than saflufenacil. In plants expressing the wild type *PPX2*, both herbicides rapidly trigger characteristic PPO inhibitor damage—intense bleaching of the youngest leaves, rapid loss of turgor, and widespread necrosis—even at the lowest doses tested. The ΔG210 allele shows the highest level of tolerance among all variants (Fig. 3). These plants maintain greener and more structurally intact rosettes across a wide dose range, especially under saflufenacil. Yet even this highly tolerant line exhibits clear, dose dependent responses to fendioxypyracil, including subtle chlorosis along leaf margins, reduced expansion of new leaves, and delayed growth to a greater extent thansaflufenacil. This contrast highlights the greater biological potency and higher activity of fendioxypyracil.

**Fig 3.**
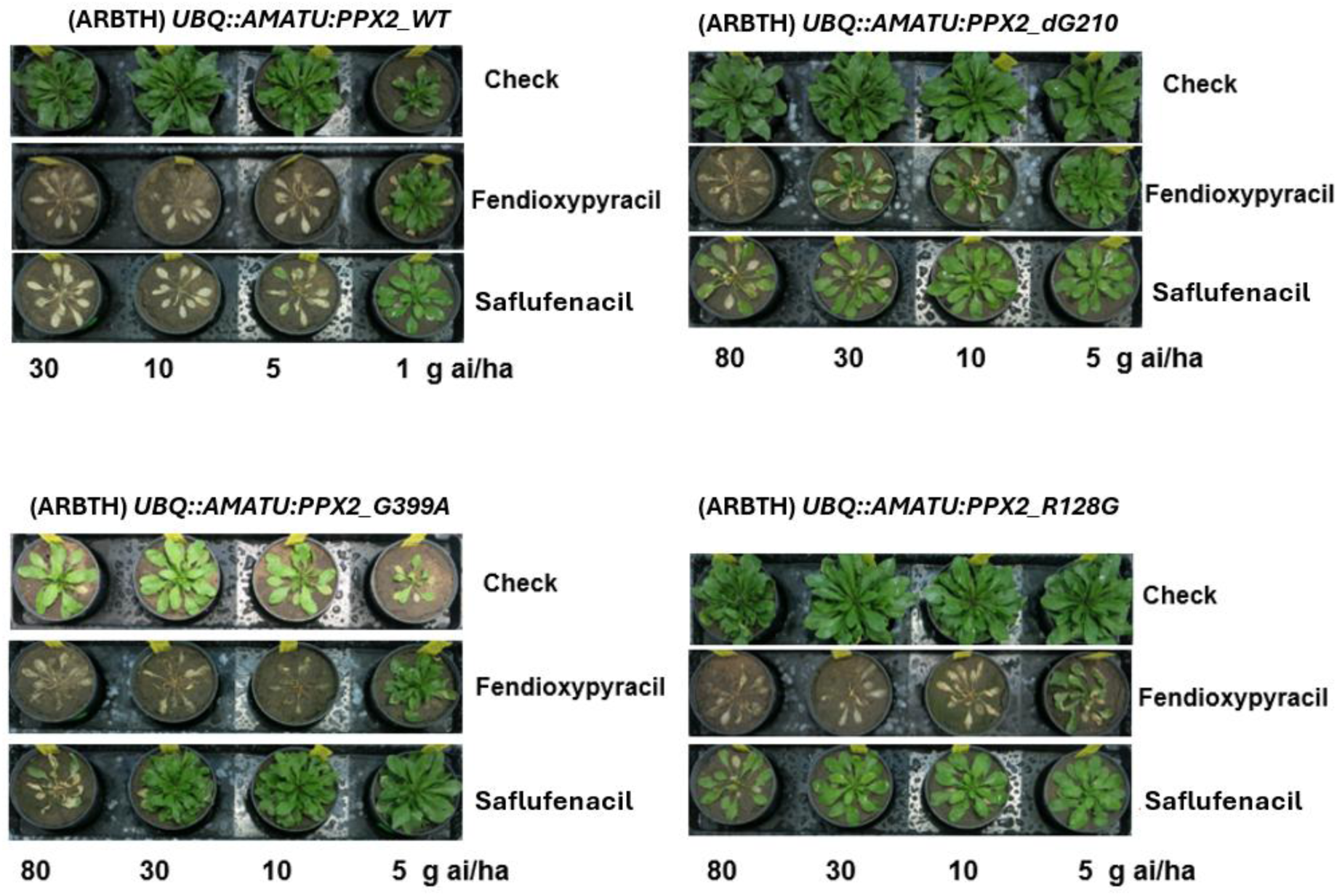
Herbicide response of Arabidopsis thaliana lines expressing different *Amaranthus tuberculatus PPX2* alleles under the UBQ promoter. Four transgenic genotypes were evaluated: *PPX2* alleles encoding wildtype, ΔG210, G399A, and R128G variants. For each genotype, the top row (“Check”) represents untreated control plants, showing normal, healthy rosettes for reference. Plants in the lower rows were treated with increasing doses of the PPO herbicides fendioxypyracil and saflufenacil.

Plants expressing G399A and R128G show high tolerance after saflufenacil treatments. Here, the difference between the two herbicides is especially pronounced: fendioxypyracil consistently causes deeper bleaching, stronger inhibition of leaf development, and earlier onset of tissue collapse, whereas saflufenacil treated plants remain relatively greener, with injury progressing more slowly and leaving portions of the rosette temporarily unaffected.

Overall, these results demonstrate that although the *Amaranthus PPX2* target site mutations confer graded levels of protection when expressed in *Arabidopsis*, fendioxypyracil is the most effective PPO inhibitor across all constructs. It produces faster, more severe, and more uniform injury than saflufenacil. This consistent pattern across all alleles underscores the superior herbicidal activity of fendioxypyracil.

### 3.4 Field evaluation of fendioxypyracil dose response

At the CCR site, herbicide significantly affected both control (χ² = 16.95, *P* = 0.009) and density reduction (χ² = 87.25, *P* < 0.001), whereas weed size (control: χ² = 0.57, *P* = 0.449; density reduction: χ² = 0.13, *P* = 0.722) and the herbicide × weed size interaction (control: χ² = 2.87, *P* = 0.826; density reduction: χ² = 10.08, *P* = 0.122) were not significant (Table 3). All fendioxypyracil rates provided control and density reduction comparable to saflufenacil and trifludimoxazin. In contrast, fomesafen resulted in significantly lower control and density reduction than the other herbicide treatments (Fig. 4,5).

**Fig 4.**
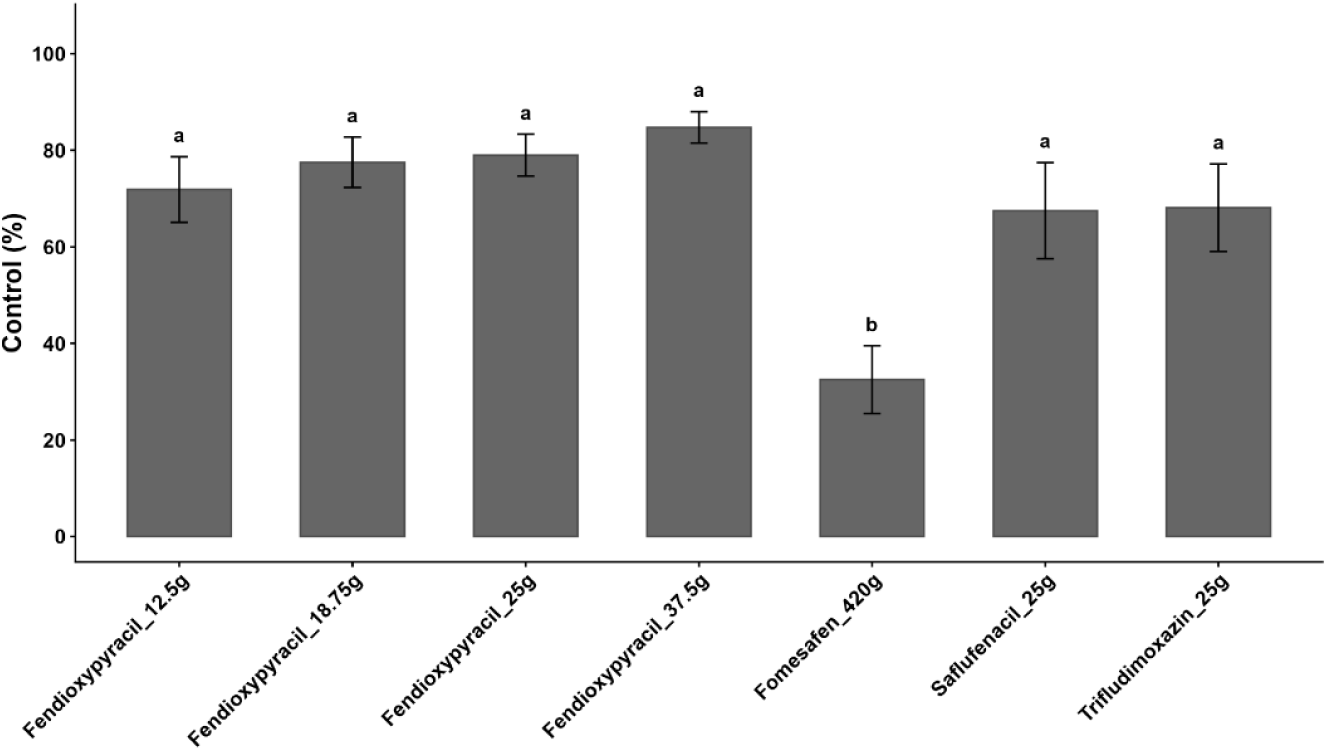
*Amaranthus palmeri* control at the CCR site. Control (%) 28 days after treatment following application of fendioxypyracil at multiple rates, trifludimoxazin (1X), fomesafen (1X), and saflufenacil (1X). Bars represent mean control, and error bars indicate standard error. Means followed by the same letter are not significantly different based on Tukey’s HSD test (P ≤ 0.05).

**Fig 5.**
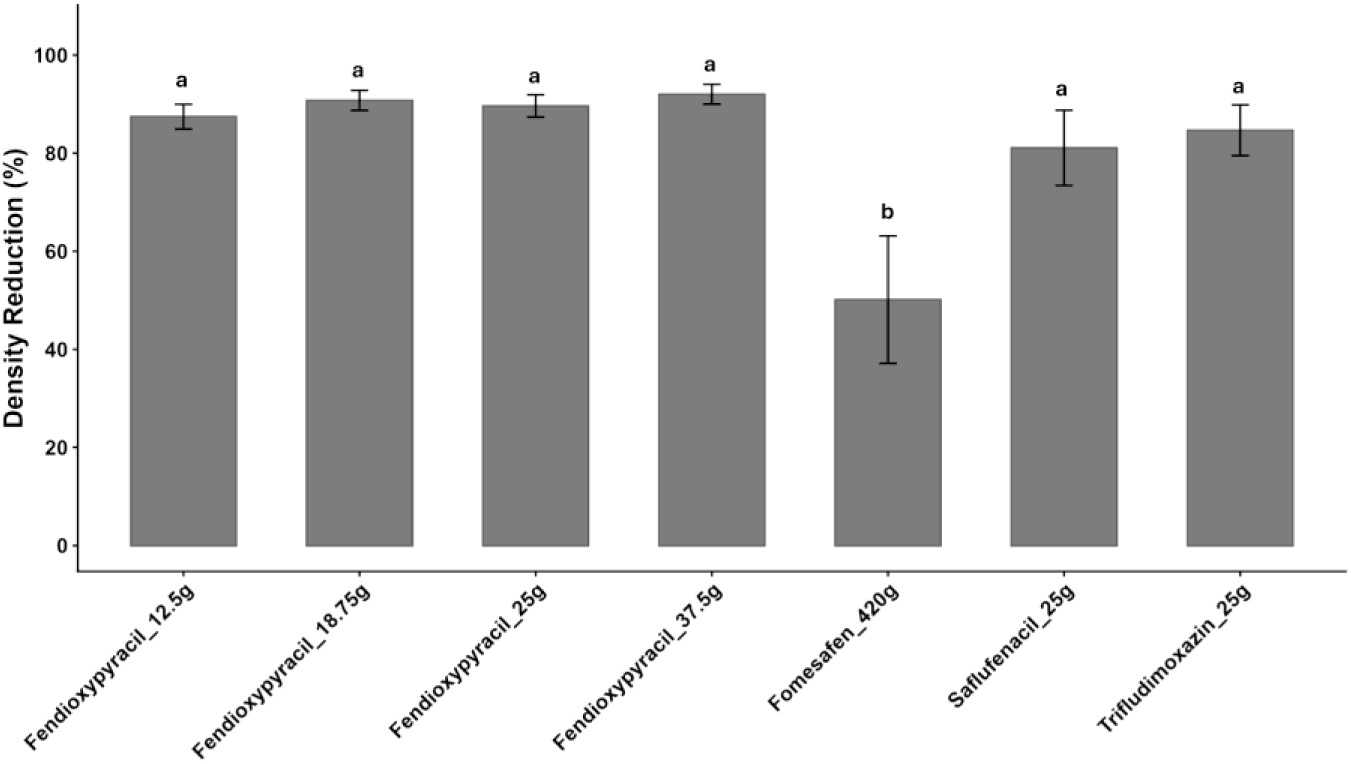
*Amaranthus palmeri* density reduction at the CCR site. Density reduction (%) 28 days after treatment following application of fendioxypyracil at multiple rates, trifludimoxazin, fomesafen (1X), and saflufenacil (1X). Bars represent mean density reduction, and error bars indicate standard error. Means followed by the same letter are not significantly different based on Tukey’s HSD test (P ≤ 0.05).

### 3.5 Prediction of future PPO resistance mutations using the *Setaria* PPO2 mutational landscape

The inhibitory activity of the PPO -inhibiting herbicides—fendioxypyracil, epyriflenacil, tiafenacil, oxadiazon, and saflufenacil was evaluated also across wild type *Setaria viridis* PPO2 and a comprehensive collection of target site mutations distributed across the three functional positions: A213, R128, and G409. These correspond to the known G210, G399 and R128 positions in *Amaranthus palmeri*, respectively. Together, these datasets provide a broad assessment of how amino acid substitutions affect herbicide potency and resistance risk in grasses. The A213 position generates the widest range of IC₅₀ values among the three panels (Table 1B).

**Table 1B.** Effect of A213 substitutions on inhibitor sensitivity of *Setaria viridis* (SETVI) PPO2. Half-maximal inhibitory concentrations (IC₅₀, M) of fendiopyracil, epyrifenacil, tiafenacil, oxadiazon, and saflufenacil are shown for wild-type PPO2 and A213 variants (A213N, A213C, A213Q, A213G, A213I, A213L, A213M, A213F, A213P, A213S, A213T, A213V). Only catalytically active variants were included in the analysis. Consequently, fewer than the 19 possible amino acid substitutions were evaluated. Values are presented in scientific notation and displayed as a heat map, where green indicates low IC₅₀ values (high sensitivity), yellow/orange indicates intermediate inhibition, and red indicates high IC₅₀ values (reduced sensitivity/resistance). Percentage remaining protein activity for each variant is indicated in the penultimate column, and the likelihood of the mutation (single vs. multiple nucleotide change) is listed in the final column.

| Enzyme | Variant (SETVI) | Fendioxypyracil | Epyrifenacil | Tiafenacil | Oxadiazon | Saflufenacil | Remaining protein activity % | Likelihood |
| --- | --- | --- | --- | --- | --- | --- | --- | --- |
| PPO2 | Wild type | 2.32E-09 | 1.81E-09 | 2.62E-09 | 1.53E-08 | 1.12E-08 | 100 |  |
| PPO2 | A213N | 4.52E-08 | 4.11E-08 | 4.66E-07 | 1.59E-05 | 6.32E-09 | 2.6 | more than 1 nucleotide change |
| PPO2 | A213C | 4.00E-09 | 1.83E-09 | 5.26E-09 | 8.26E-08 | 1.33E-07 | 100 | more than 1 nucleotide change |
| PPO2 | A213Q | 2.12E-06 | 2.56E-06 | 1.00E-05 | 1.00E-05 | 1.00E-05 | 0.6 | more than 1 nucleotide change |
| PPO2 | A213G | 8.41E-10 | 8.70E-10 | 2.48E-09 | 7.23E-08 | 9.36E-09 | 69 | 1 nucleotide change |
| PPO2 | A213I | 1.54E-09 | 3.30E-09 | 4.68E-09 | 2.65E-07 | 1.89E-07 | 3.7 | more than 1 nucleotide change |
| PPO2 | A213L | 3.03E-09 | 8.49E-09 | 2.05E-08 | 1.52E-06 | 9.51E-07 | 14 | more than 1 nucleotide change |
| PPO2 | A213M | 2.76E-08 | 1.17E-07 | 1.83E-07 | 6.23E-06 | 3.06E-06 | 8.6 | more than 1 nucleotide change |
| PPO2 | A213F | 1.25E-07 | 1.82E-07 | 5.67E-06 | 1.00E-05 | 1.00E-05 | 0.8 | more than 1 nucleotide change |
| PPO2 | A213P | 4.67E-07 | 1.80E-06 | 4.93E-06 | 1.00E-05 | 1.00E-05 | 1 | 1<br>nucleotide<br>change |
| PPO2 | A213S | 8.63E-10 | 1.22E-09 | 2.90E-09 | 4.57E-08 | 1.01E-08 | 17 | 1<br>nucleotide<br>change |
| PPO2 | A213T | 1.27E-09 | 2.16E-09 | 2.88E-09 | 4.22E-07 | 1.34E-07 | 13.7 | 1<br>nucleotide<br>change |
| PPO2 | A213V | 6.59E-10 | 1.87E-09 | 4.15E-09 | 5.69E-07 | 1.97E-07 | 11.7 | 1<br>nucleotide<br>change |

Several variants—particularly A213Q, A213P, and A213F—show dramatic reductions in herbicide potency, especially for Oxadiazon and Saflufenacil, where IC₅₀ values increase by several orders of magnitude. Even polarity changing substitutions such as A213G and A213S impair inhibitor binding to moderate degrees. Despite these effects, fendioxypyracil and epyriflenacil remain effective across nearly all A213 mutations. The capacity of these herbicides to inhibit even structurally disruptive variants highlights their robust binding mode. Only A213Q exhibits the lowest IC50s for fendioxypyracil. Hower this particular target site substitution requires more than a nucleotide change to occur, making it less likely to occur in nature. Additionally, some A213 variants show markedly reduced remaining enzyme activity, indicating that structural disruption at this position affects both enzyme activity and herbicide binding. Mutations requiring only one nucleotide change (e.g., A213G, A213V, A213S) are especially important because they might be more likely to occur, representing realistic resistance risks in field populations. Both fendioxypyracil and epyriflenacil inhibit such target site mutant enzymes.

The R128 panel displays a generally more conservative effect on herbicide potency. Most R128 substitutions—including R128G, R128I, R128L, R128S, and R128B—retain low IC₅₀ values for Fendioxypyracil, epyriflenacil, and tiafenacil (Table 1C). However, certain charged or bulky substitutions—such as R128Q, R128D, R128E, and R128W—lead to large potency losses specifically for Oxadiazon and Saflufenacil. This suggests that these herbicides rely heavily on the native cationic side chain at this location, likely due to electrostatic interactions or spatial constraints within the substrate binding pocket. In contrast, fendioxypyracil and epyriflenacil remain potent across virtually all R128 variants, reinforcing their strong mutation tolerant binding interactions. Remaining enzyme activity of the R128 variants is widely variable but often preserved to moderate levels, indicating that R128 substitutions tend to affect inhibitor binding more than catalytic stability. Many R128 mutations require only one nucleotide change, meaning that several of these altered sensitivity profiles are genetically accessible in nature. The G409 substitutions—G409A, G409E, and G409S—have surprisingly modest effects on herbicide activity. All three inhibitors—fendioxypyracil, tpyriflenacil, and tiafenacil—retain full inhibitory potency, with IC₅₀ values nearly identical to the wild type reference (Table 1D). This indicates that the structural region surrounding G409 is highly tolerant to changes without compromising inhibitor binding. In contrast, oxadiazon and saflufenacil show pronounced IC₅₀ increases, especially in G409A and G409E, suggesting that these herbicides rely on a tight steric environment that is disrupted by even minor side chain modifications. Notably, the G409 variants exhibit very low remaining enzyme activity, implying substantial structural destabilization.

**Table 1C.**
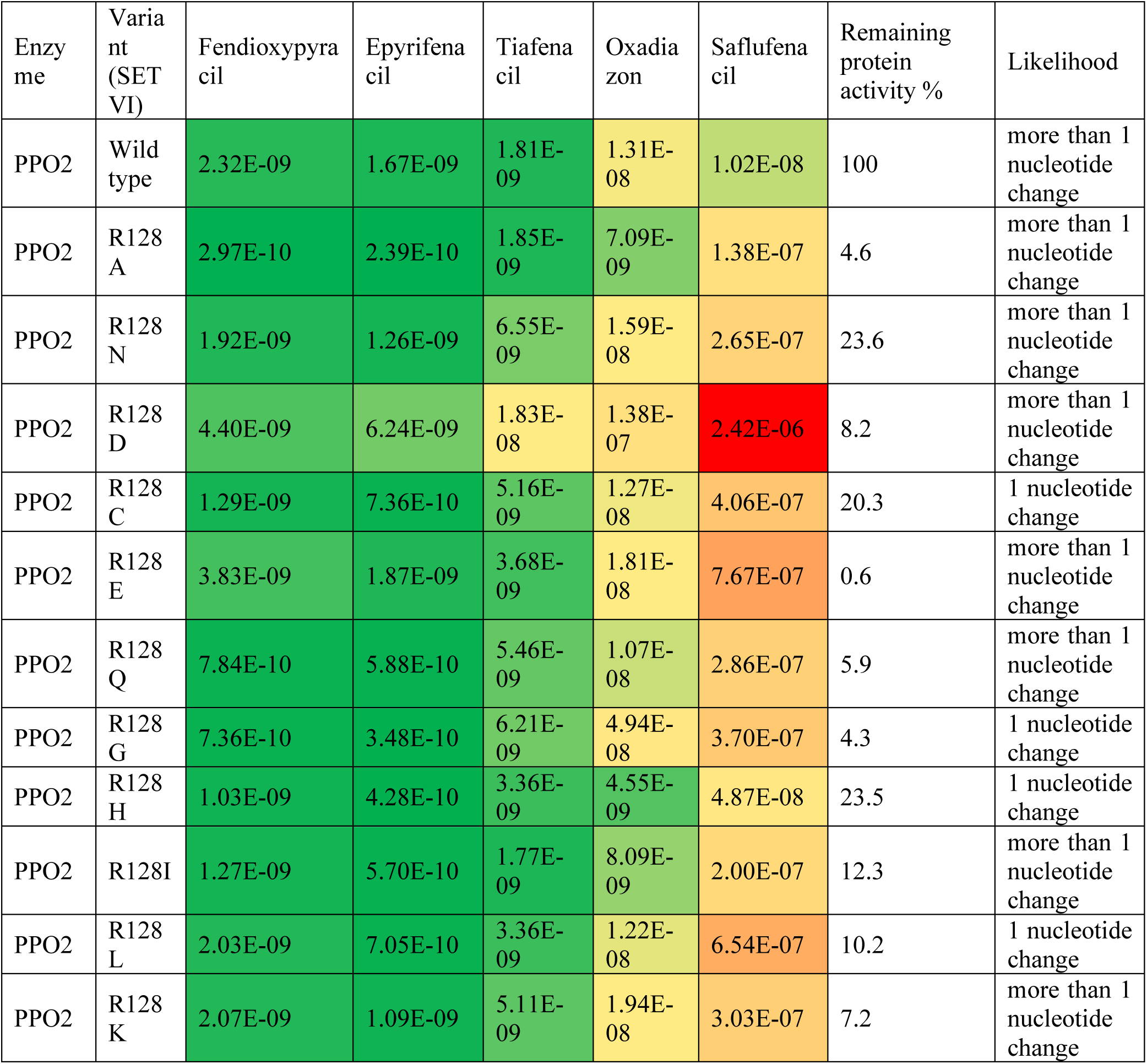
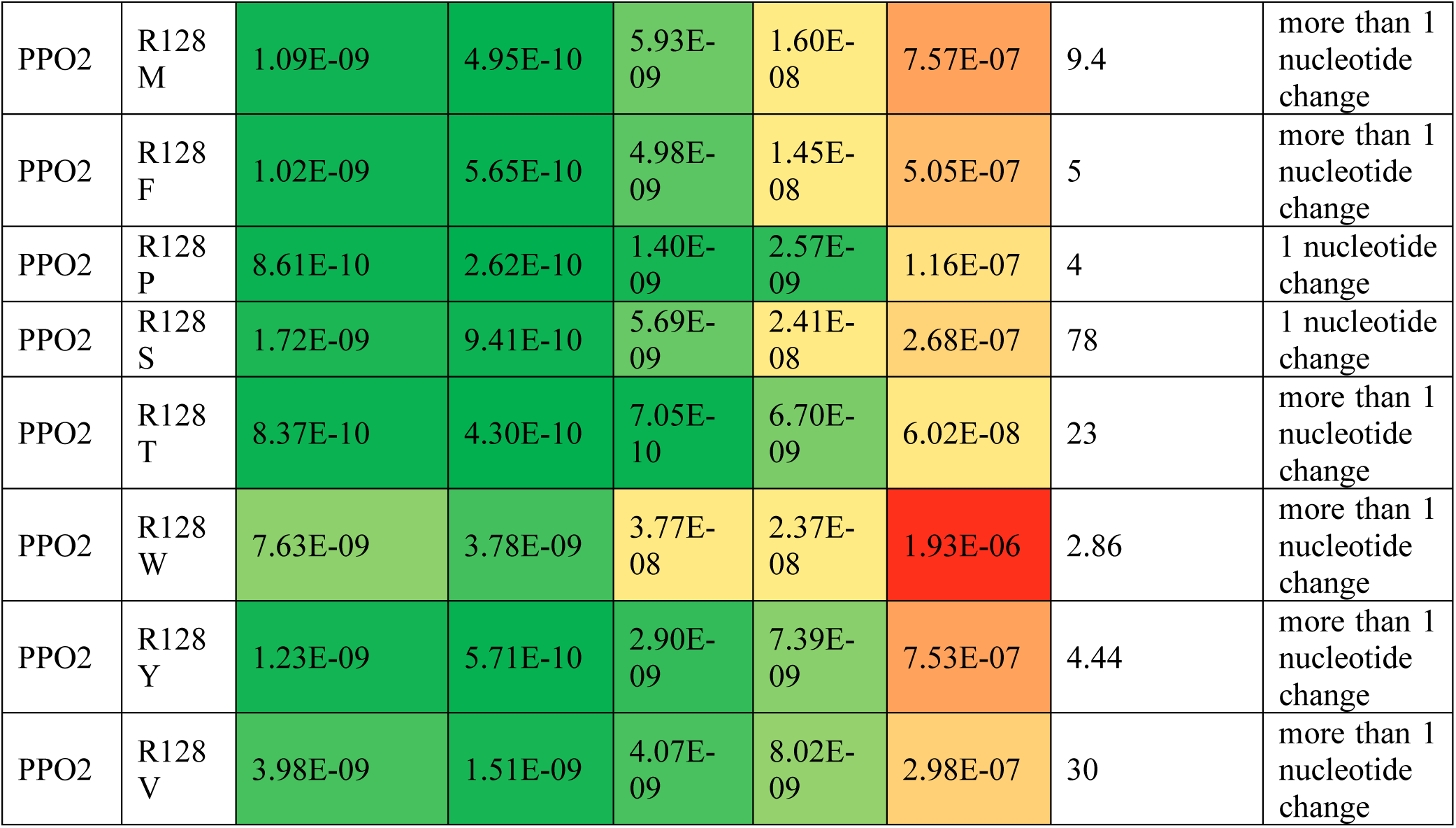
Effect of R128 substitutions on inhibitor sensitivity of *Setaria viridis* (SETVI) PPO2. half-maximal inhibitory concentrations (IC₅₀, M) of fendiopyracil, epyrifenacil, tiafenacil, oxadiazon, and saflufenacil are shown for wild-type PPO2 and a comprehensive set of R128 substitutions. Only catalytically active variants were included in the analysis. Consequently, fewer than the 19 possible amino acid substitutions were evaluated. Values are presented in scientific notation and visualized as a heat map, where green indicates low IC₅₀ values (high sensitivity), yellow/orange indicates intermediate inhibition, and red indicates high IC₅₀ values (reduced sensitivity). The percentage of remaining protein activity for each variant is reported in the penultimate column, and the likelihood of each mutation (single versus multiple nucleotide change) is indicated in the final column.

**Table 1D.**
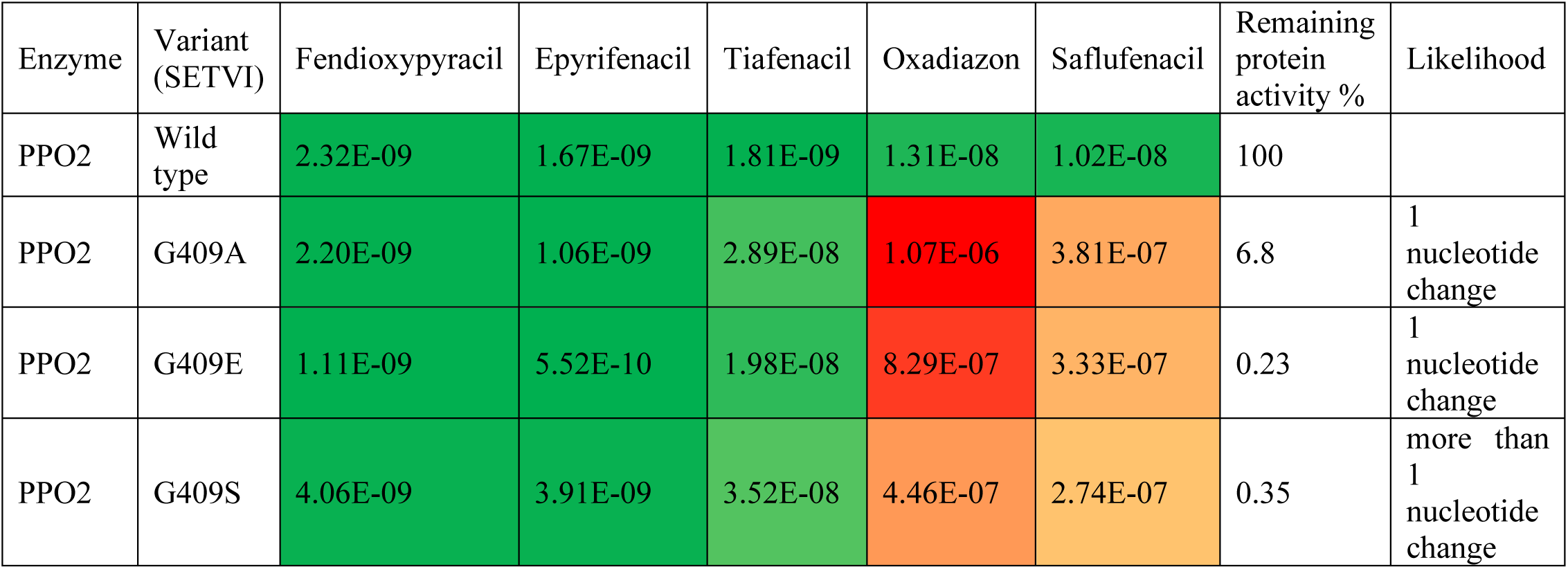
Effect of G409 substitutions on inhibitor sensitivity of *Setaria viridis* (SETVI) PPO2. Half-maximal inhibitory concentrations (IC₅₀, M) of fendiopyracil, epyrifenacil, tiafenacil, oxadiazon, and saflufenacil are shown for wild-type PPO2 and G409 substitutions (G409A, G409E, and G409S). Only catalytically active variants were included in the analysis. Consequently, fewer than the 19 possible amino acid substitutions were evaluated. Values are presented in scientific notation and visualized as a heat map, where green indicates low IC₅₀ values (high sensitivity), yellow/orange indicates intermediate inhibition, and red indicates high IC₅₀ values (reduced sensitivity). The percentage of remaining protein activity for each variant is provided in the penultimate column, and the likelihood of each mutation (single versus multiple nucleotide change) is indicated in the final column.

**Table 2:** Dose–response parameters for *Amaranthus palmeri* populations following fendioxypyracil application.

| Population | Response | Slope ($b \pm SE$) | ED<sub>90</sub> (g ai ha<sup>-1</sup> ± SE) |
| --- | --- | --- | --- |
| **S2 (Sensitive)** | GR | 3.59 ± 0.65 | < 3.00 |
| | ID | -8.69 ± 0.00 | < 3.00 |
| | LD | -4.99 ± 2.53 | < 3.00 |
| **G399A-biotype** | GR | 3.02 ± 0.20 | < 3.00 |
| | ID | -4.07 ± 1.21 | < 3.00 |
| | LD | -3.99 ± 1.27 | < 3.00 |
| **ΔG210-biotype** | GR | 1.94 ± 0.11 | 3.74 ± (0.28) |
| | ID | -1.74 ± 0.15 | 5.97 ± 0.64 |
| | LD | -1.69 ± 0.16 | 6.95 ± 0.74 |
| **R128G-biotype** | GR | 1.61 ± 0.11 | 5.0 ± (0.47) |
| | ID | -1.21 ± 0.13 | 12.10 ± 2.23 |
| | LD | -1.29 ± 0.12 | 13.23 ± 1.86 |

Dose–response relationships were fitted using a four-parameter log-logistic model. The slope parameter ( $b$ ) describes the steepness of the response curve around the inflection point. ED<sub>90</sub> represents the herbicide dose required to achieve 90% response for each measured variable. Values are expressed in g ai ha<sup>-1</sup>, where 25 g ai ha<sup>-1</sup> corresponds to the 1 × field rate. Response variables include growth reduction (GR), visible injury (ID), and mortality (LD). Values reported as “< 3.00” indicate responses below the lowest tested rate. Standard errors (SE) are provided where available.

**Table 3.** Analysis of variance (Type III Wald χ² tests) for control and density reduction at CCR site.

| Response Variable | Response | $\chi^2$ | df | P-value | Significance |
| --- | --- | --- | --- | --- | --- |
| **Control (%)** | Herbicide | 16.95 | 6 | 0.009 | ** |
|  | Weed size | 0.57 | 1 | 0.449 | ns |
| | Herbicide × Weed size | 2.87 | 6 | 0.826 | ns |
| **Density reduction (%)** | Herbicide | 87.25 | 6 | <0.001 | *** |
|  | Weed size | 0.13 | 1 | 0.722 | ns |
| | Herbicide × Weed size | 10.08 | 6 | 0.122 | ns |

Type III Wald $\chi^2$ tests were conducted using generalized linear models. Fixed effects included herbicide, weed size (*Amaranthus palmeri* size), and their interaction. ns, not significant ( $P > 0.05$ ); \* $P \leq 0.05$ ; \*\* $P \leq 0.01$ ; \*\*\* $P \leq 0.001$ .

**Table 4.** Mutation distribution of PPO2 target-site mutations in the CCR population (n = 50).

| Mutation | WT (%) | Heterozygous (%) | Homozygous mutant (%) |
| --- | --- | --- | --- |
| ΔG210 | 48 | 34 | 18 |
| R128G | 52 | 46 | 2 |
| V361A | 88 | 12 | 0 |
| G383A | 100 | 0 | 0 |
| G399A | 100 | 0 | 0 |
| F420L | 100 | 0 | 0 |

### 3.6 Can the ΔG210 target-site mutation arise in grass weeds?

Sequence alignment of the *PPX2* coding region spanning the codon encoding G210 revealed two overlapping microsatellite-like repeat motifs (**<u>TGGTGG</u>** and **<u>GTGGTG</u>**) that are present in *Amaranthus* (Fig. 6). Together, these repeats form the repetitive nucleotide sequence **TGTGGTGGA**, creating a sequence architecture predicted to facilitate DNA polymerase slippage during replication. Because the two repeat motifs partially overlap, slippage occurring within either the **TGGTGG** or **GTGGTG** repeat could result in the loss of one repeat unit. Owing to their overlapping organization, deletion of either repeat ultimately generates the same in-frame loss of a glycine codon (ΔG210), converting the encoded peptide sequence from **Cys–Gly–Gly– Asp** to **Cys–Gly–Asp** without requiring additional nucleotide substitutions.

**Fig 6.**
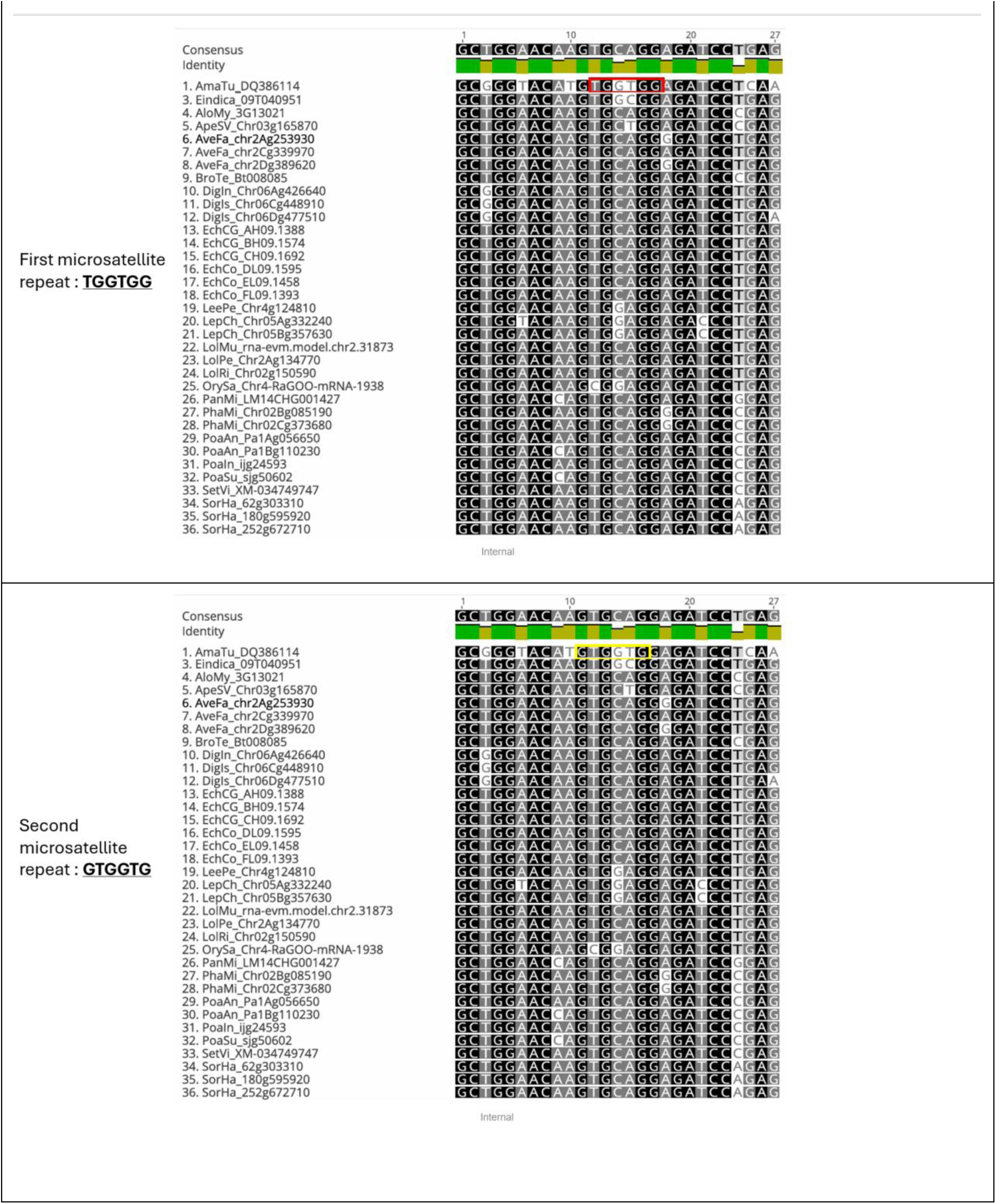
Sequence alignment of the *PPX2* DNA region encompassing two overlapping microsatellite-like repeat motifs in *Amaranthus* and representative grass species. The upper and lower panels show the first and second repeat motifs, respectively. The first motif (red box) corresponds to the tandem repeat TGGTGG, whereas the second motif (yellow box) corresponds to the overlapping repeat GTGGTG. In *Amaranthus*, these overlapping repeat architectures encode consecutive glycine residues and provide a sequence context that is consistent with DNA polymerase slippage during replication. Slippage occurring within either the TGGTGG or GTGGTG repeat motif could result in the deletion of a single TGG or GTG triplet, respectively. Because both triplets contribute to the glycine codon within the overlapping repeat architecture, either event would generate the same in-frame deletion of a glycine residue (ΔG210), shortening the encoded peptide from Cys–Gly–Gly–Asp to Cys–Gly–Asp without requiring additional nucleotide substitutions. In contrast, the corresponding *PPX2* orthologs from representative grass species (*Eleusine, Avena, Bromus, Digitaria, Lolium, Echinochloa ect.*) lack these overlapping tandem repeat motifs at the homologous position and therefore do not possess the same sequence architecture predicted to facilitate polymerase slippage. Moreover, grasses encode an alanine rather than a glycine at this position, such that an equivalent slippage event, if it occurred, would be expected to delete an alanine codon instead of a glycine codon. These alignments indicate that the overlapping repeat architecture associated with the ΔG210 deletion is conserved in *Amaranthus* but absent from the grass species analyzed.

In contrast, the corresponding *PPX2* sequences from the grass species examined (*Eleusine, Avena, Bromus, Digitaria, Lolium, Echinochloa etc.*) lack these overlapping repeat motifs and instead display non-repetitive nucleotide sequences at the homologous position (Fig. 6). Furthermore, grasses encode an alanine rather than a glycine at the equivalent amino acid position. Consequently, even if an analogous slippage event were to occur, the expected outcome would be deletion of an alanine residue rather than a glycine residue. The absence of the overlapping repeat architecture therefore suggests that the polymerase-slippage mechanism proposed to generate the ΔG210 deletion in *Amaranthus* is unlikely to operate in grass *PPX2* genes. Although alternative mutational mechanisms cannot be excluded, these sequence comparisons indicate that the repeat architecture associated with the evolution of ΔG210 is a lineage-specific feature of *Amaranthus* and is not conserved among the grass *PPX2* orthologs examined.

## 4. Discussion

The rapid spread of resistance to protoporphyrinogen oxidase (PPO)-inhibiting herbicides in *Amaranthus* spp. has become a critical challenge for weed management in row-crop systems.^4,8^ Resistance is driven by target-site mutations in *PPX2*, including the ΔG210 deletion and substitutions at R128 and G399, which collectively confer variable but often substantial reductions in herbicide sensitivity.^9,23,25^ In addition, the occurrence of mutation stacking and non-target-site resistance mechanisms has increased the complexity of resistance phenotypes and reduced the effectiveness of many established PPO inhibitors.^11–15^ In this context, the development of new PPO chemistries with improved mutation tolerance is essential to sustain the utility of this herbicide mode of action.

The present study provides a comprehensive evaluation of fendioxypyracil across multiple biological scales and demonstrates a consistent pattern of high activity against PPO-resistant systems. At the enzyme level, fendioxypyracil maintained low IC₅₀ values across a broad range of target-site mutations, including ΔG210, R128 substitutions, and G399 variants in both *Amaranthus* and *Setaria* backbone enzymes. In contrast, oxadiazon and saflufenacil showed substantial losses in potency depending on the mutation background. These results indicate that fendioxypyracil is less sensitive to structural alterations in the PPO2 binding pocket, suggesting a more robust interaction with the enzyme. This behavior is consistent with recent developments in PPO-inhibitor design, where improved binding flexibility and enhanced active-site engagement have been associated with increased tolerance to resistance mutations.^17^ Trifludimoxazin was developed to perform better against known PPO-inhibitor-resistant mutations and has shown strong enzyme-level and whole-plant activity against several common resistant mutations, including ΔG210 and other variants that reduce sensitivity to older PPO inhibitors.^17,18^ Similarly, epyrifenacil is a recently developed systemic PPO-inhibiting herbicide with broad-spectrum activity against several economically important weed species.^24^ Epyrifenacil has generally provided high levels of control across *Amaranthus* accessions, although some populations containing ΔG210 have shown reduced efficacy relative to susceptible populations.^19^. Fendioxypyracil has also shown broad-spectrum systemic activity and strong inhibition of PPO enzymes in recent studies.^21^. The low IC₅₀ values and strong whole-plant activity of fendioxypyracil across different mutation backgrounds make it a promising candidate for the management of populations carrying resistance-associated *PPX2* mutations.

Among the mutations evaluated, substitutions at position A213 in *Setaria viridis* generated the widest range of responses, with certain variants, including A213Q, A213F, and A213P, causing strong reductions in herbicide sensitivity, particularly for oxadiazon and saflufenacil. In contrast, fendioxypyracil retained high potency across most A213 variants, highlighting its ability to accommodate significant structural perturbations. Importantly, mutations that are more likely to arise under field conditions—those requiring a single nucleotide change, such as A213G, A213S, and A213V—showed only limited reductions in sensitivity while retaining substantial enzymatic activity. This combination of genetic accessibility and functional stability suggests that these substitutions may represent realistic resistance pathways; however, their limited impact on fendioxypyracil activity indicates a reduced resistance risk relative to older PPO chemistries.

The R128 position exhibited a more moderate and mutation-specific influence on herbicide sensitivity. While all substitutions had minimal effects on fendioxypyracil, certain variants, such as R128D and R128W, reduced sensitivity to oxadiazon and saflufenacil, emphasizing the importance of electrostatic interactions at this site for some herbicides. Similarly, G409 substitutions led to increased IC₅₀ values for oxadiazon and saflufenacil but had negligible effects on fendioxypyracil. Notably, many of these mutations were associated with reduced enzyme activity, indicating a trade-off between resistance and catalytic function. This trade-off is a key determinant of resistance evolution because mutations that severely compromise enzyme activity are less likely to persist in field populations.

The enzyme-level findings were strongly supported by whole-plant experiments. In *Arabidopsis* transgenic lines expressing ΔG210, G399A, and R128G PPO2 variants, fendioxypyracil consistently induced faster and more severe injury than saflufenacil across all genotypes. Even in the ΔG210 background, which is widely recognized as one of the most challenging PPO-resistance mutations,^8,16^ fendioxypyracil produced clear dose-dependent effects, whereas saflufenacil responses were weaker and less consistent. This confirms that the enhanced biochemical potency observed *in vitro* translates into improved biological activity, reinforcing the robustness of fendioxypyracil across mutation backgrounds.

Greenhouse dose–response experiments further demonstrated that fendioxypyracil maintains high efficacy against resistant biotypes. Although moderate shifts in sensitivity were observed in ΔG210 and R128G populations, effective control was achieved at the recommended field rate. In contrast, susceptible and G399A populations were controlled at rates below the lowest tested dose, indicating high intrinsic activity of the herbicide. These findings highlight an important distinction between biochemical resistance and practical field resistance, as reduced sensitivity at the enzyme level did not translate into loss of control under realistic use conditions.

Field evaluations confirmed the consistency of these results under agronomic conditions. Performance was comparable to that of trifludimoxazin and saflufenacil and exceeded that of fomesafen, particularly at larger weed sizes, where older PPO inhibitors typically show reduced efficacy. The absence of major efficacy losses across sites indicates that fendioxypyracil can maintain performance across diverse genetic backgrounds and environmental conditions.

Although fendioxypyracil performed well under field conditions, some limitations should be considered. Only a single resistant population and field environment were evaluated; therefore, additional testing across a broader range of *Amaranthus palmeri* populations and environmental conditions is needed to determine the consistency of these responses. Fendioxypyracil provided significantly greater control and density reduction than fomesafen while maintaining efficacy comparable or better to newer PPO chemistries, suggesting improved activity against resistance-associated PPO2 variation. Genotype data for the CCR population, including PPO2 target-site mutations, are provided in Table 4.

An additional important aspect highlighted in this study is the species-specific nature of certain resistance mechanisms. The ΔG210 deletion, a dominant resistance mechanism in *Amaranthus*, arises from a replication-slippage event facilitated by a tandem-repeat motif that is absent in grass *PPX2* sequences.^8,23^ This structural feature suggests that some resistance pathways may be inherently limited to specific taxonomic groups, reducing the likelihood of their occurrence in grass weeds. Such insights are relevant for predicting resistance evolution and assessing cross-species risk.

Despite the strong performance of fendioxypyracil observed in this study, long-term sustainability will depend on appropriate resistance-management strategies. The extensive history of PPO-inhibitor use demonstrates that repeated selection pressure can lead to the accumulation of target-site mutations, mutation stacking, and the evolution of metabolic resistance.^4,11–15^ Therefore, the integration of fendioxypyracil into diversified weed-management programs, including herbicide rotation and mixtures with alternative modes of action, will be essential to preserve its effectiveness.

In summary, fendioxypyracil represents a next-generation PPO inhibitor with a broad and robust activity profile across major target-site resistance mutations. Its consistent performance at the enzyme, plant, and field levels demonstrates reduced sensitivity to structural alterations in PPO2 and highlights its potential as a valuable tool for managing PPO-resistant *Amaranthus* populations.

## Acknowledgements

The authors acknowledge BASF for funding and thank all co-authors for their contributions. Open AI was used to improve language and grammar of the manuscript.

## Conflict of interest statement

The authors declare no conflicts of interest.

## Notes

### Competing Interest Statement

The authors have declared no competing interest.

